# Longitudinal change in executive function is associated with impaired top-down frontolimbic regulation during reappraisal in older adults

**DOI:** 10.1101/867689

**Authors:** William K. Lloyd, Jayne Morriss, Birthe Macdonald, Karin Joanknecht, Julie Nihouarn, Carien M. van Reekum

## Abstract

Networks in the prefrontal cortex (PFC) that are important for executive function are also engaged in adaptive responding to negative events. These networks are particularly vulnerable to age-related structural atrophy and an associated loss of executive function, yet existing evidence suggests preserved emotion processing ability in aging. Using longitudinally acquired data from a battery of cognitive tasks, we defined a metric for the rate of decline of executive function. With this metric, we investigated relationships between changes in executive function and emotion reappraisal ability and brain structure, in older adults using functional and structural MRI. During task-based fMRI, participants were asked to cognitively reappraise negatively valenced images. We hypothesised one of two associations with decreasing executive function over time: 1) a decreased ability to reappraise reflected in decreased PFC and increased amygdala activation, or 2) a neural compensation mechanism characterised by increased PFC activation but no differential amygdala activation. Structurally, for a decreased reappraisal ability, we predicted a decrease in grey matter in PFC and/or a decrease of white matter integrity in amygdala-PFC pathways. Neither of the two hypotheses relating to brain function were completely supported, with the findings indicating a steeper decline in executive function associated with both increased PFC and increased left amygdala activity when reappraising negative stimuli. In addition, white matter integrity of the uncinate fasciculus, a primary white matter tract connecting the amygdala and ventromedial areas of PFC, was lower in those individuals who demonstrated a greater decrease in executive function. These findings highlight that when cognitive ability diminishes, active top-down emotional control is compromised.

## 1. Introduction

Ageing is characterised by marked neural changes in the brain, with direct consequences for mental functioning and wellbeing. These changes include a spatially heterogeneous loss of grey matter and a reduction in white matter integrity, where age-related atrophy is more pronounced in areas such as the medial temporal lobe and prefrontal cortex (Giorgio et al. 2010; Gunning-Dixon et al. 2009). Such age-related neural changes have been associated with impairments in memory and “executive function” (e.g. selective attention, switching, inhibition of irrelevant information; Buckner, 2004; Raz & Rodrigue, 2006; Van Petten et al., 2004). These brain regions susceptible to age-related atrophy overlap with brain networks supporting successful regulation of emotion (for a review, see Mather, 2012).

The ability to adaptively respond to negative situations is crucial for maintaining mental and physical health. Impaired regulation of emotion due to brain atrophy could render older adults particularly vulnerable to mood disorders, such as depression and anxiety. Given the overlap in neural networks involved in executive function and emotion regulation, and patterns of age-related brain atrophy, the notion that executive function and emotion regulation are interlinked and should both decline with age is compelling (e.g. Urry & Gross, 2010). Yet, behavioural studies suggest improved emotion regulatory function with advancing age, possibly due to a reduced need for cognitive effort to achieve a regulatory goal in older age (Scheibe and Carstensen 2010). A number of factors have been suggested to contribute to this apparent paradox, including the use of different regulation strategies with advancing age (Mather 2012). Faced with structural brain changes and a decline in executive function, how do older adults maintain emotion regulatory function?

### The emotion regulation network

Animal and human research (Banks et al. 2007; Carmichael and Price 1995; Ongur and Price 2000) has demonstrated bidirectional anatomical connections between the amygdala (a subcortical structure that plays a key role in the acquisition and expression of emotion) and medial and orbital areas of the prefrontal cortex (PFC) (which are important for higher order cognition and flexible value assignment respectively). The connection between these regions is considered instrumental in adaptive responding to emotional situations (Nomura et al., 2004, Banks et al. 2007). Using a paradigm of voluntary emotion regulation, human neuroimaging studies have identified a larger distributed network of prefrontal-amygdala brain regions subserving emotion regulation (Beauregard et al. 2001; Eippert et al. 2007; Goldin et al. 2008; Kalisch et al. 2006; Kim and Hamann 2007; Kevin N. Ochsner et al. 2002; K. N. Ochsner et al. 2004; Phan et al. 2005; Heather L Urry et al. 2006; van Reekum et al. 2007). These regions include the amygdala and ventromedial PFC as mentioned, but also involve lateral and dorsomedial PFC (DMPFC) (for a recent meta-analysis see Morawetz, Bode, Derntl, & Heekeren, 2017). Given the role of lateral PFC in maintaining representations and adjusting implementations, and DMPFC in self-referential processing and performance monitoring, these areas are thought to regulate negative emotion through cognitive reconstrual, or “reappraisal” of the emotion-eliciting stimulus (e.g. Braunstein et al., 2017).

Most prior studies have identified increased PFC and decreased amygdala activation when participants reappraise negative stimuli with the goal to reduce negative affect, with a key role for the ventrolateral prefrontal cortex (VLPFC) in reappraisal success (Wager, Davidson, Hughes, Lindquist, & Ochsner, 2008). Furthermore, work from our group has demonstrated that the ventromedial PFC (VMPFC) mediates the inverse association between left VLPFC and amygdala in non-depressed adults (Johnstone et al. 2007) and between DMPFC and amygdala in healthy older adults (Urry et al., 2006) when reappraising negative stimuli.

### Emotion regulation in the ageing brain

A few recent studies have examined functioning of this emotion regulatory network in healthy younger and older adults. Opitz et al. (2012) report lower left VLPFC and DMPFC regulation-related activation in older compared to younger adults, but no differential age effect in amygdala activation. Using a similar voluntary emotion regulation task, Winecoff et al. (2011) have also demonstrated lower activation for older compared to younger adults in left VLPFC but, again, no age difference in subcortical regions such as the amygdala. However, the same study showed that age-independent higher cognitive function, as measured using a composite score based on tasks designed to assess memory and executive function, is associated with a greater decrease in amygdala activation. In sum, these findings demonstrate that emotion regulation-associated PFC activation changes with age, and cognitive ability is an important indicator of emotion regulation outcome and is reflected in amygdala activity.

### Structural and functional changes in the ageing brain

A common view is that age-related changes in brain function are caused, at least in part, by structural changes in the brain. This view is supported by evidence relating structural brain changes to behavioural outcomes (Morcom and Johnson 2015; Park and Reuter-Lorenz 2009; Winecoff et al. 2011). Yet, only few studies have directly examined relationships between brain structure, function and cognitive decline. While some recent reports suggest a relationship between grey matter atrophy and cognitive decline (Fleischman et al. 2014; Manard et al. 2016), the overall body of evidence has been weak (Salthouse 2011). Studies incorporating measures of grey matter density and cognitive brain function in older adults reveal conflicting results. One study has found higher activation associated with higher grey matter density (Teipel et al., 2007), while other studies have shown higher activation associated with *lower* grey matter density (Kalpouzos, Persson, & Nyberg, 2012). A further study found a positive association between grey matter volume and activation, but no age association (Burgmans et al. 2011). Our work on emotional processing in older age found a compensatory “shift” in engagement from lateral to medial PFC regions that was associated with grey matter loss in lateral PFC and age (van Reekum et al., 2018).

Other research suggests that variability in white matter, rather than grey matter, is more likely coupled to cognitive functioning in older adults, with stronger associations between executive function scores and white matter integrity than cortical thickness (Ziegler et al., 2010). A recent longitudinal study extended these findings by demonstrating increased activity in lateral frontal cortex during task switching associated with a decline in white matter integrity in tracts connecting PFC regions (Hakun, Zhu, Brown, Johnson, & Gold, 2015).

In summary, multimodal brain imaging studies that integrate structural brain changes and the effects of such changes on brain function are sparse, and hitherto non-existent in the field of emotion regulation.

### The overlap between emotion regulation and executive function

Even at the behavioural level, only recently have researchers begun to examine executive function performance alongside emotion regulation in ageing (e.g. Gyurak, Goodkind, Kramer, Miller, & Levenson, 2011; Gyurak et al., 2009). These findings suggest that aspects of executive function, such as verbal fluency, predict emotion regulatory ability in older adults. Yet, other research points at a general association between “fluid” cognitive abilities, including working memory and processing speed, and emotion regulation, in the absence of an effect of age group (Opitz, Lee, Gross, & Urry, 2014). This is despite older adults showing lower fluid cognitive abilities compared to younger adults.

The reliance on comparison between younger and older adult groups, where between-group difference is assumed to be a proxy measure of ageing and/or cognitive decline, has hampered the majority of past research on emotion regulation and ageing. We argue that cohort effects or a lifetime’s experience with emotional situations may be confounded with age. Furthermore, such between-group analyses will not adequately account for variation in, or preservation of, cognitive ability in the older age group. Therefore, we propose to examine associations between cognitive ageing and emotion regulation by focusing on change in executive function *within* a middle-to-older age group, taking advantage of longitudinal behavioural data to measure the rate of decline of cognitive ability for each participant.

### Aims and predictions

The aim of the present study is to test the relationship between executive function, brain structure, and emotion regulation-related brain function in older adults, by examining associations between a longitudinal measure of executive function and functional and structural MRI. We hypothesized that the rate of change of longitudinal executive function (rLEF), over and above any age or cross-sectional EF performance differences, would predict activity in regions of the brain instrumentally associated with emotion regulation (VMPFC, VLPFC and amygdala specifically), as well as structural brain integrity in this network. Specifically, we hypothesized one of two functional associations with decreasing longitudinal executive function: either a decreased ability in emotion downregulation reflected in decreased prefrontal cortical (PFC) activation and increased amygdala activation, or a compensatory effect with increased PFC activation but no differential amygdala effect (cf. van Reekum et al., 2018). Structurally, we predicted a decrease in grey and/or white matter integrity of amygdala-prefrontal cortical pathways that would mediate the functional response of the amygdala and PFC.

## 2. Materials and Methods

### 2.1. Participants

34 older-aged participants (aged 56-83, M age = 73.1, SD = 6.8, 20 (57.1%) female) were included in this study. This group was derived from a larger cohort of 70 older adults recruited from the Reading University Older Adult Research Panel, via local newspaper and poster advertisements, and 20 younger adults also recruited via advertisements. We included all older adult participants for whom we had longitudinal cognitive data; that being at least two measures of executive function spaced at least two years apart. All participants scored between 24-30 on the Mini-Mental State Examination (MMSE, see Table 1 for details on standard neuropsychological test performance at the time of testing). No participants reported a history of neurological disorders or current use of steroid medication. All participants were right-handed. Participants were also screened for contraindication to MR scanning. This study was carried out in accordance with the declaration of Helsinki (1991, p. 1194). All procedures were approved by the NHS Research Ethics Service and the University of Reading’s Research Ethics Committee. Participants provided written informed consent prior to their participation.

**Table 1.** Descriptive statistics from sample and cognitive tasks

|  | N | Minimum | Maximum | Mean | Std. Deviation |
| --- | --- | --- | --- | --- | --- |
| Age | 34 | 56 | 83 | 72.90 | 6.90 |
| MMSE_total | 34 | 24 | 30 | 27.50 | 1.50 |
| CERAD_total | 34 | 2 | 3 | 2.37 | 0.29 |
| PI_Mod | 32 | -60.67 | 370.34 | 104.08 | 117.25 |
| Stop (ssd) | 31 | 108.50 | 859.60 | 493.02 | 192.48 |
| Stroop (i/c) | 30 | 0.50 | 1.20 | 1.08 | 0.13 |
| Trails B/A | 34 | 1.20 | 3.90 | 2.20 | 0.73 |
| Verbal Fluency -<br>Letter | 34 | 10 | 26 | 17.53 | 4.26 |
| Verbal Fluency -<br>Category | 34 | 11 | 31 | 21.62 | 5.54 |

### 2.2. Stimuli and tasks

#### 2.2.1. Stimuli

96 images were selected from the International Affective Picture System (Lang et al. 2008). Images were selected based on the normative valence ratings and categorised as either negative (76 images, valence = 1.46-3.85, M = 2.41, SD = 0.57) or neutral (24 images, valence = 4.38-5.88, M = 5.10, SD = 0.36). Picture categories were matched on luminosity, complexity and social content.

#### 2.2.2. Emotion regulation task

The voluntary emotion regulation task comprised the reinterpretation of negative events, such as scenes depicted in images or movie clips, to lessen or intensify an emotional response. Similar forms of the task have been previously used with studies involving older adults (Opitz et al. 2012; Heather L. Urry et al. 2009). To induce negative affect prior to regulation, participants viewed each picture for three seconds after which they were presented with an audio instruction to “suppress” (decrease), “enhance” (increase), or “maintain” (attend). Upon hearing “suppress” participants were asked to consider a less negative outcome of the scenario than they had thought prior to the instruction. For “enhance” they were asked to consider a more negative outcome, and for “maintain” they were asked to keep their original impression in mind. Participants were asked to keep following the instruction until the image disappeared after a further six seconds. Participants received training and practice on the task immediately prior to scanning^1^. Example regulation strategies were suggested, e.g. for a picture of an injured animal to “enhance” their response they might consider the animal will die, or to “suppress” they might consider the animal will recover. Participants then practiced the task on five scenarios and were asked what strategy they used to follow the instruction. Training and practice was repeated if participants did not report the use of emotion regulatory strategies as intended.

Following the picture, a ratings screen was presented for two seconds, during which participants were asked to rate the prior image as neutral, somewhat negative, quite negative, or very negative. The 96 trials involved a pseudo-randomised presentation of picture-instruction combinations. All neutral pictures were accompanied by the “maintain” instruction.

The protocol was divided into four blocks, between which the participant was allowed to rest before commencing the next block. Inter trial interval was jittered, and each block lasted for a duration of approximately seven minutes.

#### 2.2.3. Stimulus presentation

Stimuli were coded and presented using E-Prime (version 2.0.10.242, Psychology Software Tools, Inc., Pittsburgh, PA, USA) and delivered via VisualSystem binoculars (VisualSystem, NordicNeuroLab, Bergen, Norway). The VisualSystem binoculars were also used to monitor the participant’s eye movements (eye movement data were not further analysed) and state of alertness throughout the experiment. Response to the ratings screen was recorded using a four-button MR-compatible button box held in the participant’s right hand.

#### 2.2.4. Cognitive tasks

As part of the Older Adult Research Panel, participants are invited to perform a battery of cognitive tests upon intake, and again every two years thereafter. Where participants had not been (re)tested within two years prior to the scan date, they were asked to resit the test battery as part of their participation of this study. Tests included, amongst others, verbal fluency (Miller, 1984) and trail making tests (Reitan & Wolfson, 1985). The verbal fluency test included a letter and a semantic/category component. Participants were asked to name out loud as many words as possible in one minute starting with the letter F and then name as many animals as possible in one minute. The session was recorded and the experimenter counted the number of correct words, ignoring duplicates. For the trail making test, participants were tasked with drawing a line as quickly as possible to connect the elements in a sequence, initially with one category (numbers) and then with two alternating categories (numbers and letters). The time for both tasks was recorded and scores were obtained by dividing time (number and letters) / time (numbers).

Other tests that were part of the Older Adult Research Panel test battery were collected but were not included in the composite score due to a lack of compatible historical data (e.g. switching from paper-and-pencil based tasks to computerised tasks, such as for the Stroop task), and are not reported here.

### 2.3 Data reduction and analysis

#### 2.3.1. Executive function data reduction

Longitudinal data from verbal fluency category^2^ and trail-making tasks, was used to calculate a longitudinal measure of the rate of change of executive function (rLEF) for each older adult, whereby the most negative rLEF score indicates the steepest decline in performance. For each participant, each set of data points included information of time since initial test in years (with the first test being at time point zero). A linear regression was used to calculate slope (i.e. the rate of change). Of the 35 participants, 11 had two timepoints, 14 had three, and 10 had four. The mean slope of verbal fluency was 0.16 (std dev = 1.01) and trail making was −0.03 (std dev. = 0.144) (see supplementary material for further details). To assess specificity for longitudinal, rather than cross-sectional EF performance, we equally computed a composite score across the raw scores of these two tasks acquired in the last session prior to scanning.

#### 2.3.2. MRI data acquisition

MRI data was collected using a 3T Siemens Magnetom scanner and 12-channel head coil (Siemens Healthcare, Erlangen, Germany) at the University of Reading Functional Imaging Facility. A 3D-structral MRI was acquired for each participant using a T1-weighted sequence (Magnetization Prepared Rapid Acquisition Gradient Echo (MPRAGE), repetition time (TR) = 2020 ms, echo time (TE) = 3.02 ms, inversion time (TI) = 900 ms, flip angle = 9°, field of view (FOV) = 250 x 250 x 192 mm, resolution = 1 mm isotropic, acceleration factor = 2, averages = 2, acquisition time = 9 min 7 s). Emotion regulation task-related fMRI data was collected in four identical blocks, using an echo planar imaging (EPI) sequence (211 whole-brain volumes, 30 sagittal slices, slice thickness = 3.0 mm, slice gap = 33%, TR = 2000 ms, TE = 30 ms, flip angle = 90°, FOV = 192 x 192 mm^2^, resolution = 3 mm isotropic, acquisition time = 7 min 10 s per block). Diffusion-weighted images (DWI) were collected with an EPI sequence (50 sagittal slices, slice thickness = 2.0 mm, TR = 6800 ms, TE = 93 ms, resolution = 2 mm isotropic, averages =2, acquisition time = 7 min 24 s). Diffusion weighting was distributed along 30 directions with a *b* value of 1000 s/mm^2^. For each diffusion set, one volume with a *b* value of 0 s/mm^2^ was also acquired. Further structural and functional MRI data were acquired as part of this protocol but are not reported here.

#### 2.3.3. FMRI preprocessing and registration

Preprocessing and statistical analysis of MR data was carried out in FSL 5.0.7 (FMRIB’s Software Library, www.fmrib.ox.ac.uk; Jenkinson, Beckmann, Behrens, Woolrich, & Smith, 2012; Smith et al., 2004). Subject FMRI data were field-map corrected to adjust for distortions due to magnetic field inhomogeneity, spatially realigned to correct for head motion, slice-time corrected, normalized to MNI space via coregistration to the subject T1-weighted image, and smoothed using a Gaussian 6 mm FWHM kernel.

#### 2.3.4. FMRI Analyses

Data were modelled at block level with three explanatory variables (EVs) used to describe the instruction conditions (“suppress”, “enhance”, “maintain”) with negative pictures, and one EV for the “maintain” instruction with neutral pictures. A further EV, used to model the valence judgement, and six motion correction EVs were included as covariates of no interest. At the intermediate level, mean effect for each condition was determined in a fixed effects model across the four blocks. At the final level, GLM analysis consisted of regressors for rLEF, cross-sectional EF, and age^3^ using FSL’s Randomise with threshold free cluster enhancement estimated from 5000 permutation samples per independent test. All significant statistical analyses are reported for p < 0.05, familywise error (FWE) corrected.

Pre-defined anatomical masks were created for bilateral amygdala and VLPFC (Harvard-Oxford atlas, Desikan et al., 2006), as well as VMPFC (McGill, Mazziotta, Toga, Evans, Fox, & Lancaster, 1995) using a 50% probability threshold. These masks were used for small volume correction of the voxel-wise search in those regions. Contrasts of negative/suppress vs negative/maintain (suppress vs maintain), and negative/enhance vs negative/maintain (enhance vs maintain) were assessed with age and rLEF as regressors. Results are reported with peak MNI coordinates, number of voxels, and max Z.

#### 2.3.5. Grey matter probability (GMP)

Similar to the methods reported in van Reekum et al. (2018), we used FSL’s FAST algorithm (Zhang et al., 2001) to segment the high resolution T1-weighted image into grey matter, white matter, and CSF. We then normalised and blurred the GMP maps to the same extent as the functional data of interest. Cluster-wise GMP was extracted using the statistical ROIs from the FMRI regression analysis and associations between GMP and functional data (residualised for age and cross-sectional EF performance, per the performed voxel-wise search with rLEF), were assessed using Pearson correlations, with an alpha level of *p* < 0.05.

#### 2.3.6. DWI preprocessing, registration, and analyses

Diffusion weighted data were corrected for subject head motion and the effects of eddy currents. Fractional anisotropy (FA) maps were produced by fitting calculated tensor values to the data. Analysis of FA data was performed using FSL’s tract-based spatial statistics (TBSS) method. This involves the removal of the most distal voxels to remove likely outliers, nonlinear registration of subject data to a study-specific template and affine alignment to the MNI152 T1-weighted template. All subject data were used to determine the mean white matter FA skeleton, which was thresholded at a factor of 0.2. A mask of this skeleton was created to define the extent of the voxelwise analysis of FA data. Anatomical masks of the left and right uncinate fasciculus, forceps minor and corticospinal tract were used for region of interest analysis. These masks were taken from the JHU-ICBM-DTI probabilistic atlas (Mori, Wakana, Zijl, & Nagae-Poetscher, 2005) thresholded at 0.25. The forceps minor and corticospinal tract served as control regions, similar to previous structure-function research (Swartz, Carrasco, Wiggins, Thomason, & Monk, 2014). We used rLEF as a predictor variable of white matter integrity as measured by tract-based fractional anisotropy. Age and cross-sectional EF were included in the model as covariates. Statistical analyses of FA data were performed using FSL’s Randomise with threshold-free cluster enhancement estimated from 5000 permutation samples per independent test. All significant statistical analyses are reported for *p* < 0.05, familywise error (FWE) corrected.

## 3. Results

### 3.1.1 Effects of longitudinal change in executive function on brain activation in an emotion regulation task

Associations between rLEF and contrasts of activation were assessed. We focused our analyses on a-priori regions of interest in PFC (VLPFC and VMPFC) and amygdala, based on their association with emotion regulation. For completeness, whole-brain results outside of these a-priori ROIs are also reported (see Tables 2 and 3).

**Table 2.** A-priori PFC and amygdala regional activation patterns in response to stimuli presented in the emotion regulation task

| Contrast | Brain region | Voxels | Max Z | Location of max Z |  |  |
| --- | --- | --- | --- | --- | --- | --- |
|  |  |  |  | x | y | z |
| Suppress > Passive | L VLPFC | 1058 | 6.18 | -42 | 14 | 24 |
| Suppress > Passive | R VLPFC | 76 | 4.28 | 56 | 32 | -2 |
| Suppress > Passive | R VLPFC | 15 | 2.36 | 46 | 20 | 10 |
| Suppress > Passive | R VLPFC | 8 | 4.06 | 58 | 28 | 16 |
| Suppress > Passive x rLEF | L VLPFC | 114 | 4.11 | -50 | 8 | 18 |
| Suppress > Passive x rLEF | R VLPFC | 2 | 3.59 | 52 | 28 | 8 |
| Suppress > Passive x rLEF | R VLPFC | 1 | 3.6 | 56 | 30 | 10 |
| Suppress > Passive x rLEF | VMPFC | 17 | 3.9 | -4 | 48 | 8 |
| Suppress > Passive x rLEF | VMPFC | 16 | 4.32 | 8 | 36 | 2 |
| Suppress > Passive x rLEF | L Amygdala | 54 | 4.51 | -22 | -8 | -14 |
| Suppress > Passive x rLEF | R Amygdala | 43 | 4.71 | 24 | -2 | -16 |
| Enhance > Passive | L VLPFC | 1303 | 5.97 | -48 | 20 | 26 |
| Enhance > Passive | R VLPFC | 1013 | 5.95 | 48 | 32 | -2 |
| Enhance > Passive | VMPFC | 218 | 3.81 | -8 | 48 | -6 |
| Enhance > Passive | R Amygdala | 30 | 3.58 | 30 | 2 | -20 |
| Enhance > Passive | L Amygdala | 2 | 3.65 | -24 | -2 | -20 |

Note: Corrected cluster for multiple comparisons at $p < 0.05$ . Location of cluster's maximum Z are in MNI space. R = right; L = left. rLEF = the rate of change of longitudinal executive function.

**Table 3.** Whole-brain activation patterns in response to stimuli presented in the emotion regulation task

| Contrast | Brain region | Voxels | Max Z | Location of max Z |  |  |
| --- | --- | --- | --- | --- | --- | --- |
|  |  |  |  | x | y | z |
| Suppress > Passive | Bilateral Superior Temporal Gyrus, extending into Bilateral Hippocampus, Bilateral Caudate, Bilateral Insula, **Bilateral VLPFC**, Bilateral Middle Frontal Gyrus, Superior Frontal Gyrus, Paracingulate, Cingulate Left Angular Gyrus | 22174 | 7.23 | 54 | -20 | -6 |
| Suppress > Passive | R Angular Gyrus, R Middle Temporal Gyrus | 895 | 5.54 | 46 | -58 | 50 |
| Suppress > Passive | R Precentral Gyrus | 37 | 3.77 | 20 | -24 | 66 |
| Suppress > Passive | L Pallidum | 25 | 4.17 | -14 | -14 | -10 |
| Suppress > Passive | L Thalamus | 14 | 3.13 | -10 | -16 | -4 |
| Suppress > Passive | R Postcentral Gyrus | 9 | 2.85 | 18 | -34 | 74 |
| Suppress > Passive | R Precentral Gyrus | 3 | 3.05 | 10 | -26 | 64 |
| Suppress > Passive | L Postcentral Gyrus | 2 | 2.9 | -4 | -38 | 76 |
| Suppress > Passive x rLEF | Bilateral Frontal Operculum Cortex, extending into Bilateral Central Opercular Cortex, **Bilateral Amygdala**, Bilateral Insula, Bilateral Pre/Post Central Gyrus, **Bilateral VLPFC**, Cingulate, Paracingulate, **VMPFC** | 18425 | 5.66 | 36 | 22 | 4 |
| Suppress > Passive x rLEF | Precuneus Cortex, Intracalcarine Cortex, Lingual Gyrus | 181 | 4.73 | 8 | -60 | 6 |
| Suppress > Passive x rLEF | R Postcentral Gyrus | 38 | 2.84 | 42 | -32 | 48 |
| Suppress > Passive x rLEF | L Heschl's Gyrus | 36 | 3.46 | -50 | -28 | 6 |
| Suppress > Passive x rLEF | R Amygdala | 36 | 3.01 | 30 | 6 | -26 |
| Suppress > Passive x rLEF | R Parahippocampal Gyrus | 14 | 3.18 | 16 | -30 | -16 |
| Suppress > Passive x rLEF | L Middle Frontal Gyrus | 12 | 3.81 | -24 | 2 | 50 |
| Suppress > Passive x rLEF | R Frontal Pole | 8 | 3.28 | 34 | 34 | 18 |
| Suppress > Passive x rLEF | Cingulate Gyrus | 7 | 2.67 | 4 | -42 | 24 |
| Suppress > Passive x rLEF | L Middle Frontal Gyrus | 6 | 3.9 | -32 | 10 | 56 |
| Suppress > Passive x rLEF | R Middle Frontal Gyrus | 6 | 3.02 | 54 | 16 | 40 |
| Suppress > Passive x rLEF | L Frontal Pole | 6 | 3.17 | -32 | 60 | 2 |
| Suppress > Passive x rLEF | Lingual Gyrus | 6 | 3.75 | -2 | -66 | 2 |
| Suppress > Passive x rLEF | R Superior Frontal Gyrus | 4 | 2.78 | 24 | -8 | 66 |
| Suppress > Passive x rLEF | L Frontal Orbital Cortex | 4 | 2.7 | -56 | 30 | -10 |
| Suppress > Passive x rLEF | R Middle Frontal Gyrus | 3 | 2.64 | 40 | 14 | 38 |
| Enhance > Passive | Bilateral Angular Gyrus, extending into Bilateral Occipital Cortex, **Bilateral Amygdala**, Bilateral Hippocampus, Bilateral Thalamus, Bilateral Caudate, Bilateral Insula, Bilateral Central Opercular, Bilateral Pre/Post Central Gyrus, **Bilateral VLPFC**, Cingulate, Paracingulate, **VMPFC** | 135926 | 7.66 | 50 | -48 | 16 |

Note: Corrected cluster for multiple comparisons at $p < 0.05$ . Location of cluster's maximum Z are in MNI space. R = right; L = left. rLEF = the rate of change of longitudinal executive function. Bold font denotes whole-brain activation falling within a-priori regions of interest.

To test for either a decreased ability in emotion regulation (reflected in decreased PFC and increased amygdala activation during reappraisal), or a compensatory effect (increased PFC activation but no change in amygdala), we first regressed rLEF against the activation contrast of suppress vs maintain. rLEF correlated negatively with the suppress vs maintain activation contrast in both PFC regions of interest; VMPFC, and left VLPFC, as well as bilateral amygdala (see Figures 1, 2 and Table 2). In other words, greater activation in these regions to suppress vs maintain was associated with a higher rate of EF decline. In the whole brain analysis, negative correlations between rLEF and the suppress vs maintain activation contrast were also observed in the bilateral insula, bilateral post-/pre-central gyrus, bilateral intracalerine cortex/precuneus, posterior cingulate gyrus and paracingulate/anterior cingulate gyrus, among others (see Table 2). There were no significant correlations of rLEF with the enhance vs maintain contrast in a-priori regions, nor in the whole brain analysis.

**Figure 1.**
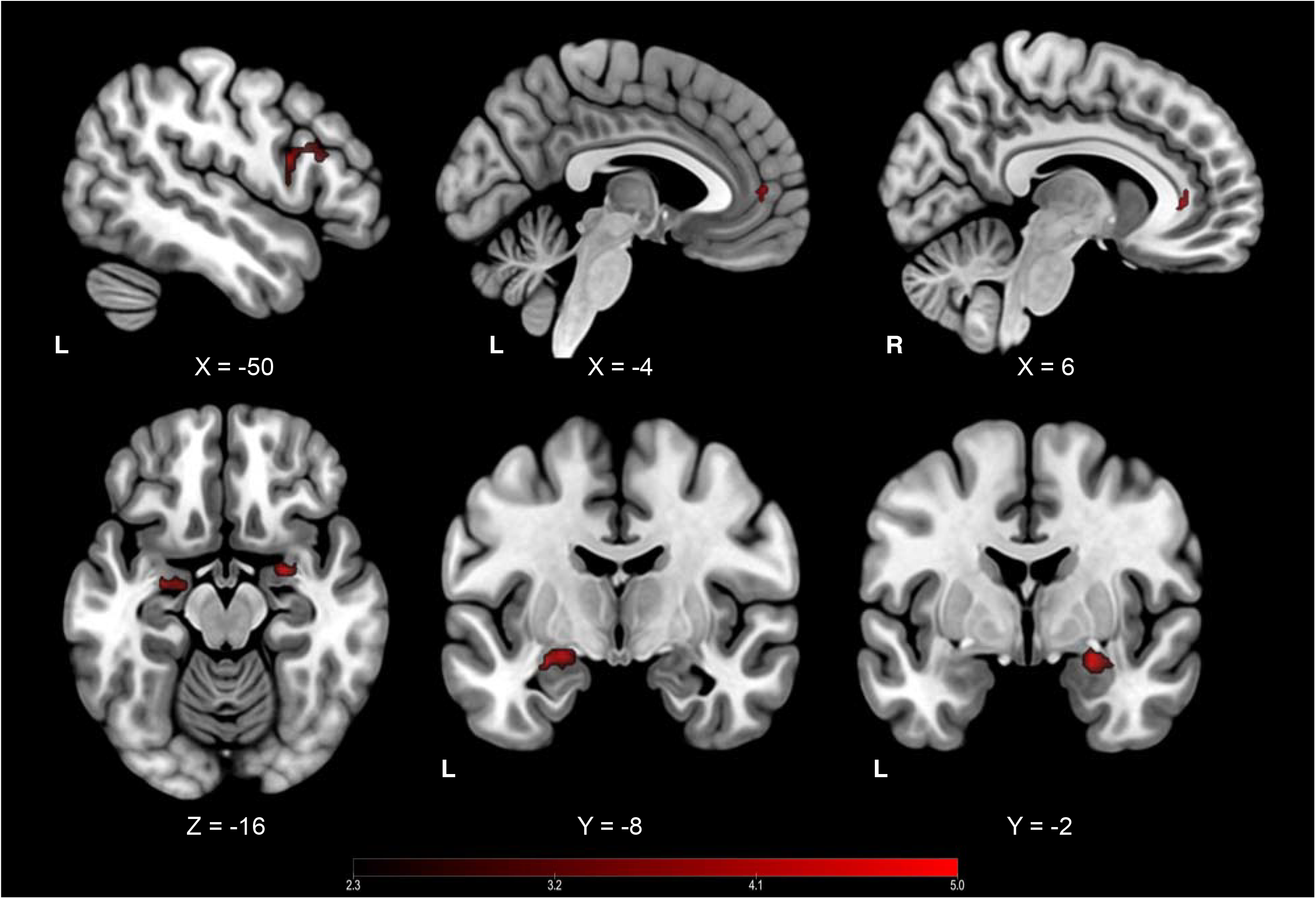
Significant correlations for the contrast of suppress vs maintain associated with rLEF, at ROI (red) corrected level of significance thresholding. Coordinates in MNI space: L, left; R, Right. rLEF = the rate of change of longitudinal executive function.

**Figure 2:**
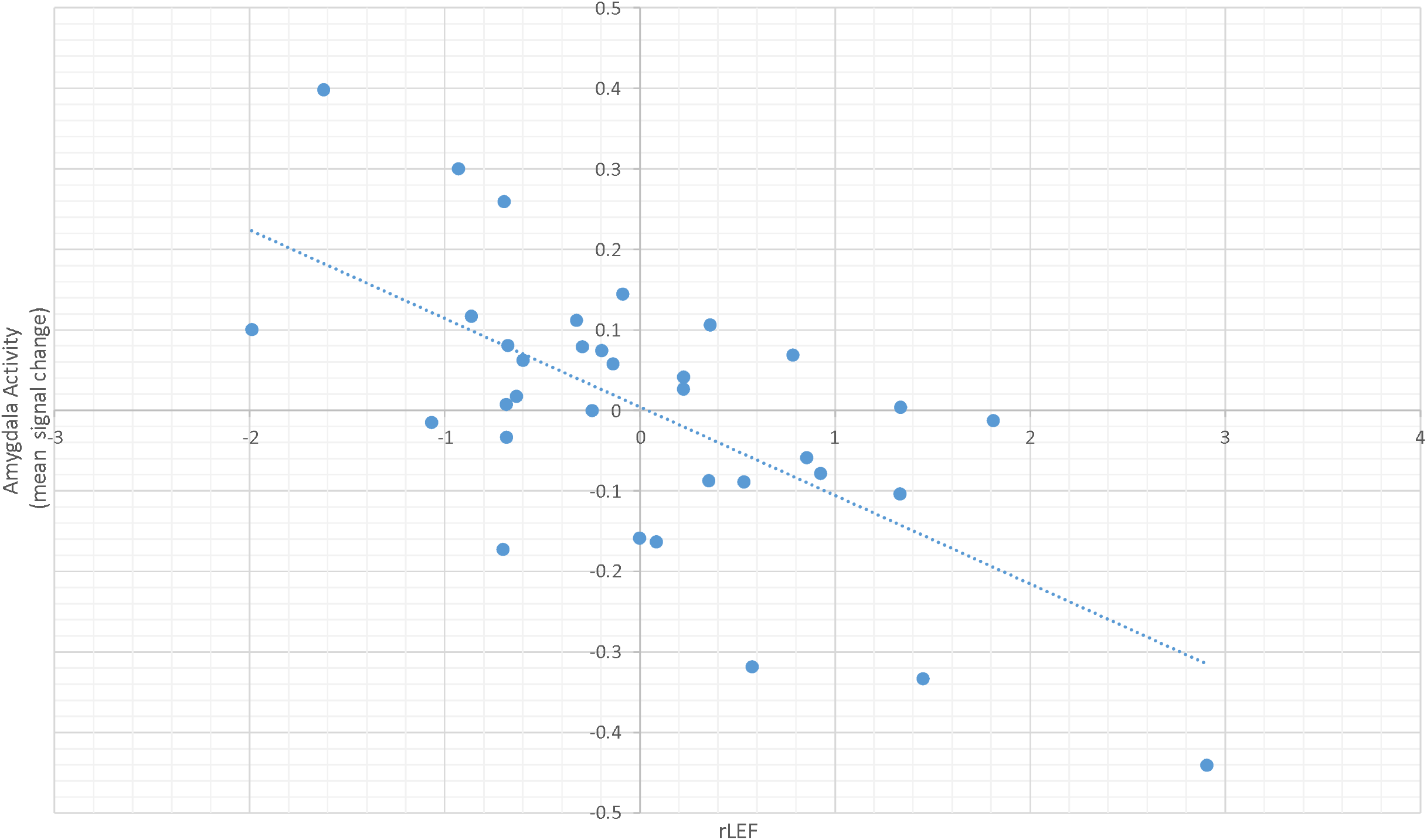
Graph demonstrating the inverse linear relationship between rLEF and amygdala activation (residualised for cross-sectional EF and age) for the suppress v maintain contrast.

To ensure that these patterns were unique to rLEF, we calculated variance inflation factors (VIFs) for rLEF, cross-sectional EF (i.e. measured around the time of scanning) and age. All VIFs were sufficiently low (age=1.08, EF=1.32, rLEF=1.29) to allow the assumption that multicollinearity can be safely ignored. Relatedly, there were neither ROI nor whole brain associations with age or cross-sectional cognitive performance for the suppress vs maintain contrast or for the enhance vs maintain contrast.

### 3.1.2 Main effects

For main effects only, we found typical patterns of activation for the emotion regulation task in the a-priori regions of interest for suppress vs maintain (increased activation in bilateral VLPFC) and enhance vs maintain contrasts (increased activation in bilateral amygdala, bilateral VLPFC and VMPFC), and in the whole brain analysis (see Tables 2 and 3). We did not find a main effect in amygdala for maintain vs. suppress.

#### 3.2. Grey and white matter integrity and associations with BOLD responses

Results from the task-based FMRI experiment demonstrated emotion regulation-based changes in activation associated with rLEF in, among others, left VLPFC, VMPFC and left amygdala. We assessed associations between the BOLD response in the PFC clusters and the GMP within and across these areas, to test the prediction that rLEF-related differences in BOLD response patterning would be associated with individual differences in PFC structural integrity (see van Reekum et al., 2018 for findings suggesting some functional compensation for age-related grey matter loss). While there was a trend towards a negative correlation between the GMP in VMPFC and the BOLD response in left VLPFC for the suppress relative to the maintain condition, this correlation was not significant, *r*(32) = -.29, *p* = .10. None of the other correlations were significant (max *r*(32) = .20, all others *r* < .1). GMP in these prefrontal regions was also not correlated with rLEF, max *r* = .16, *p* = .36.

We next assessed the integrity of the uncinate fasciculus, the primary white matter tract that connects the VMPFC and amygdala. We used rLEF as a predictor variable of white matter integrity as measured by tract-based fractional anisotropy. Age and cross-sectional EF ability were included in the model as covariates. Using the pre-defined left and right uncinate fasciculus masks as regions of interest, results showed a significant positive correlation between FA and rLEF in the left uncinate fasciculus (number of voxels = 40, p < 0.01, MNI coordinates: −30,8,-8), situated between the insula and putamen. A smaller region was also seen in right uncinate (number of voxels = 5, p < 0.07, MNI coordinates: 2,31,10) (See Figure 3). FA values in the control regions (forceps minor and corticospinal tract) did not show any clusters of voxels significantly correlated with rLEF. Thus, greater structural integrity of the uncinate fasciculus was specifically associated with maintenance of executive function over time.

**Figure 3.**
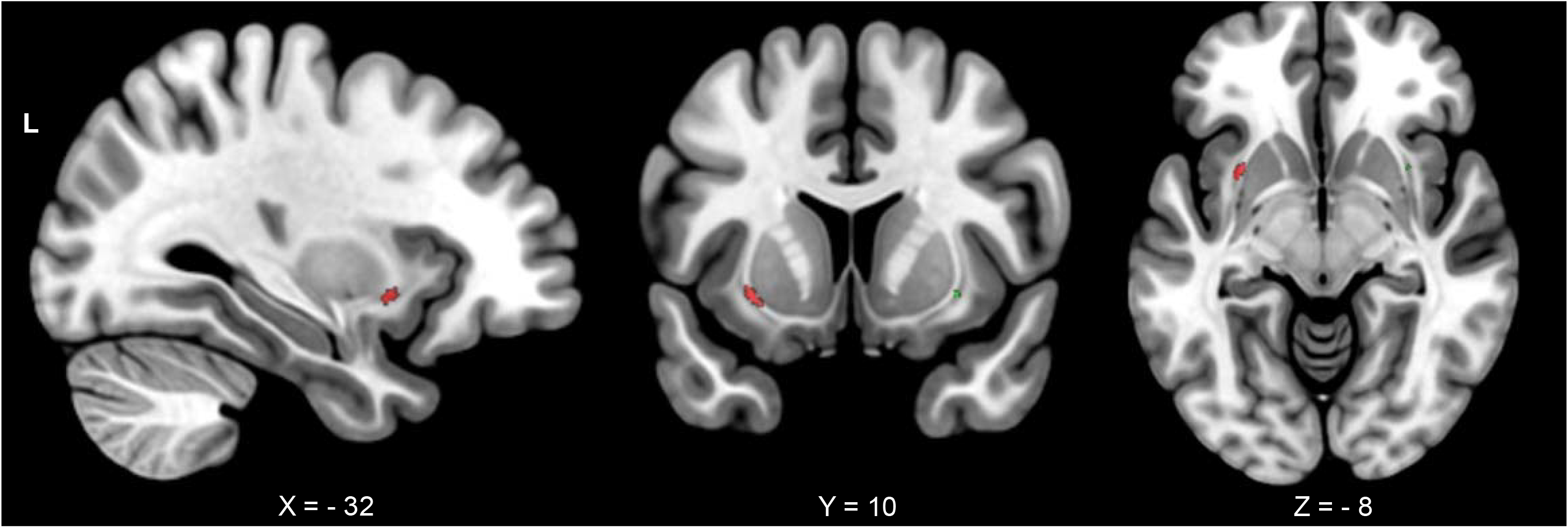
Significant negative correlations for rLEF with FA in the left (red) and right (green) uncinate fasiculus, controlling for age. Coordinates in MNI space; L, left. rLEF = the rate of change of longitudinal executive function.

The BOLD response in bilateral amygdala for the suppress relative to attend condition showed a trend towards a negative correlation with FA in the left uncinate fasciculus cluster, r(31) = -.33, p = .062, such that higher uncinate fasciculus integrity in this region was associated with a stronger reduction in amygdala when asked to decrease the emotional response. Furthermore, rLEF significantly mediated the association between FA in this cluster in the uncinate fasciculus tract and amygdala activation (Indirect effect size = −0.76, SE = 0.32, 95% CI = [-1.33, −0.11]. However, FA did not mediate the amygdala-rLEF association (Indirect effect size = 0.00, SE = 0.02, 95% CI = [-0.04, 0.02]). Mediation analyses were performed using PROCESS (Hayes (2018) www.guilford.com/p/hayes3) implemented in SPSS v20.

#### 3.3. Are the ratings predicted by rate of change in EF?

During the emotion regulation task, after each image, participants were asked to score the negativity of the image on a four-point scale, with 1 reflecting “neutral” and 4 “very negative”. Reflecting the FMRI analytic approach, we assessed whether the negativity ratings for the pictures presented in the task differed as a function of regulation (enhance, attend, suppress) and rLEF. We also took age and cross-sectional EF ability (all continuous predictor variables were demeaned) into consideration in the MANCOVA model. The rate of change did not explain variance in the ratings, over and above age and cross-sectional EF performance, *F* = .67, *p* = .52, *n.s.*. A main effect of regulation was found on unpleasantness ratings, *F*(2, 29) = 12.26, *p* <.001. On average, rated negativity was higher for the “enhance” condition (*M*=2.75, *SD*=0.40) compared to the “maintain” condition (*M*=2.51, *SD*=0.34). Unexpectedly, the “suppress” condition was associated with higher rated negativity of the images (*M*=2.60, *SD*=.37) compared to the maintain condition. With respect to cross-sectional EF and age, age interacted with regulation, *F*(2, 29) = 3.55, *p* = .042. Follow-up comparisons to further investigate the age x regulation interaction was carried out by estimating the ratings means at +1 and −1 SD of the mean age respectively. Results suggested that “older” older adults provided more differential ratings of the images than the “younger” older adults as a function of regulation condition, with both the “enhance” and “suppress” conditions resulting in higher negative ratings (*M* = 2.73 for enhance and *M* = 2.55 for suppress) than the “maintain” conditions (*M* = 2.36, *p* < .001 and *p* = .006 resp.), and the negative ratings being higher in the enhance vs suppress condition (*p* = .024). This effect was not significant for the “younger” group (at −1 SD, with the multivariate *F*(2,29)=1.74, *p*=.192; *M* enhance = 2.78, *M* maintain = 2.65, *M* suppress = 2.66).

## 4. Discussion

Our findings demonstrate that, when decreasing negative emotion through reappraisal, activation in the fronto-limbic network is greater in individuals with a larger negative rate of change of executive function. More specifically, we found that a relative decline in EF predicted reduced downregulation of activity in left amygdala, despite recruitment of various PFC regions that are key in emotion regulation, including VMPFC and VLPFC (e.g. Morawetz et al., 2017). Outside these a-priori regions of interest, a decline in EF was also associated with recruitment of other areas in (pre)frontal cortex including frontal pole, anterior cingulate and premotor and supplementary motor cortex, and areas in insula, pre/post central gyrus, posterior cingulate cortex, and precuneus. The association between the rate of change in EF and activation in these brain regions was specific to the decrease condition, as no significant associations were observed in the increase vs maintain contrast. Focusing on individual differences in integrity of the uncinate fasciculus, a white matter tract connecting amygdala with prefrontal cortical areas including VMPFC, we showed that a higher rate of decline of executive function was associated with decreased white matter integrity in left uncinate fasciculus and, at trend-level, also in right uncinate fasciculus. Grey matter probability in VMPFC and VLPFC areas demonstrating activation differences as a function of rLEF was not significantly associated with BOLD response, nor with rLEF.

Prior brain imaging work on reappraisal-based emotion regulation has presented relatively few age-related differences in neural activity. Across two studies, reduced recruitment of left VLPFC was found for older compared to younger adults (Opitz et al., 2014, Winecoff et al., 2011) but these studies found no age-related differences in other brain areas, including the amygdala. Furthermore, increased cognitive performance, including EF, has been shown to predict greater regulation-related decreases in amygdala activation (Winecoff et al. 2011) and reappraisal success (modelled using a composite of self-report, physiological and expressive behaviour measurements, Opitz et al., 2014) *regardless* of age. Our findings add to this knowledge base to suggest that, in older age, reappraisal is compromised not necessarily when age is high or EF is low, but when a given person’s EF ability decreases over time.

In line with this suggestion, we found that the white matter integrity of a cluster in the left uncinate fasciculus, a primary pathway between VMPFC and amygdala, decreased with increasing rate of cognitive loss. This finding was specific to the uncinate fasciculus; no such correlations were observed in the control tracts. It may be the case that the loss of structural integrity in this tract means that the successful recruitment of key areas in the prefrontal cortex is not translated into a suppression of amygdala activity, although the direct relationship between higher uncinate fasciculus integrity and decreased bilateral amygdala activity when participants were reappraising the negative information showed only a trend-level correlation. There was however, a full mediation effect of longitudinal EF on this association, suggesting that longitudinal EF change carried the relationship between reappraisal-related amygdala activity and white matter integrity. Given that the clusters included in the mediation analysis were derived from a voxel-wise regression analysis with rLEF as the predictor variable, and that the association between amygdala and white matter integrity in the uncinate fasciculus was not strong, this is perhaps not surprising. Future research on structure-function relationships in ageing would benefit from data-driven approaches, such as those discussed by Calhoun (2018), to further address how age-related structural loss affects brain function. Unlike prior findings suggesting functional compensation for age-related grey matter loss in left VLPFC during picture-induced negative affect (van Reekum et al., 2018), we did not find any associations with grey matter probability and either rLEF or BOLD response in PFC regions in this study. Whilst interpreting a null finding is tricky, it could be that variability in grey matter in these PFC regions is more tightly coupled to age per se, and less to cognitive functioning in older age, per recent studies (Ziegler et al., 2010; Hakun et al., 2015).

Longitudinal change in EF was not only positively associated with the BOLD response to the suppress condition in a-priori prefrontal areas but demonstrated associations in a larger network of regions, including anterior cingulate cortex and insula, typically involved in salience/attentional control (see e.g. Seeley 2007). Connectivity between salience and arousal networks has been shown to increase with heightened arousal (Young et al., 2017). We found a decline in EF related to increased engagement of these brain regions when decreasing negative affect. Prior work has demonstrated that cognitive ageing is often paired with increased activation and disrupted connectivity in similar cortical regions, often thought to underlie a compensation for loss of brain structure integrity (for a review, see Sala-Llonch et al., 2015). Our findings add to this literature by highlighting that it is not age per se, nor group-relative EF performance per se, but an age-related change in EF that may engage such compensatory mechanisms. In other words, an over recruitment occurs when individuals no longer function at the level that they were accustomed to. The specificity of the findings to the suppress (and not the enhance) condition further suggests that this condition is marked by individual differences specifically, a point made earlier by our group (Urry et al., 2006) and others (for a review see Ochsner et al., 2012).

With respect to the ratings of unpleasantness as a function of regulatory instruction, we found that the pictures were rated as more negative after both regulation instructions compared to the attend condition. The ratings did not vary as a function of longitudinal change in EF, over and above age and cross-sectional EF. We did find that older age within our sample was associated with higher ratings of picture unpleasantness, both after the instruction to increase and decrease their response to the pictures. Higher negative ratings after the instruction to decrease emotion have previously been reported in research involving older adults (e.g. Opitz et al. 2012) and the reason for this effect is not entirely clear. The effect could be due to different, and in general more, visual scanning of the pictures after receiving an instruction to regulate to generate a narrative serving reappraisal, interacting with the ratings provided of the picture, rather than the experience. Prior work involving older adults has identified that the instruction to reappraise with the goal to increase or decrease picture-induced negative affect was associated with a higher number of fixations and larger distances between fixations (van Reekum et al., 2007). Increased scanning may have resulted in an increased intake of information, enhancing the negativity rating of the picture, irrespective of their experience of negative affect. Indeed, whilst we found on average increased left and right amygdala activation for the enhance condition, no such effect was found for the suppress condition (see Table 2). This finding has previously been reported, particularly in studies involving older adults and where instructions were delivered during, rather than prior to, picture presentation (e.g. Urry et al. 2006, 2009). The reason for asking about the rating of the picture rather than their experience was to counter any potential demand characteristics resulting from providing an explicit instruction. Future work should assess other, ideally implicit, indicators of emotional experience.

Given the relatively sparse literature that examines both executive function and emotion regulation in the same older adult cohort, this work provides an important addition to the body of evidence on how executive function and emotion regulation are coupled. As stated, some behavioural evidence suggests that markers of executive function are positively associated with emotion regulation ability in cross-sectional samples of older adults (e.g. Gyurak, Goodkind, Kramer, Miller, & Levenson, 2011; Gyurak et al., 2009). We sought to expand on this evidence in two ways; firstly by focusing on a longitudinal measure of executive function to obtain each individual’s rate of decline, and secondly by correlating this metric with structural and functional brain patterns supporting emotion regulation. We found that individuals showing the largest rate of decline also show the largest activation in amygdala and PFC when regulating emotion. These findings may suggest that a preservation of emotion regulation ability with age until executive function starts to decline. What remains to be explained, however, is why increased PFC activation was not paired with a decrease in amygdala activation in our study. Whilst requiring replication, our findings provide evidence of a more complex scenario whereby decline in white matter integrity inhibits successful regulation despite increased functional effort.

In conclusion, the findings presented here highlight that the rate of change of executive function, and not age or cognitive ability per se, impacts the ability to downregulate picture-induced negative affect despite increased cortical engagement. With respect to structure-function integration, findings suggest that the integrity of key white matter tracts supporting corticolimbic interaction, rather than grey matter atrophy, may impede downregulation of amygdala responsivity when the context requires one to do so. These findings have implications for our understanding of age-related cognitive decline and may be relevant for pathology associated with rapid cognitive decline (e.g. Alzheimer’s, dementia). The ability to adaptively respond to negative situations is crucial for maintaining mental and physical health. Impaired regulation of emotion due to brain atrophy could render older adults particularly vulnerable to mood disorders, such as depression and anxiety. Further longitudinal studies are warranted to map the “tipping point” in age-related cognitive decline and emotional well-being.

## Supplementary Material

### Rate of change of longitudinal executive function (rLEF)

rLEF was calculated by combining scores from verbal fluency category and trail-making tasks, collected longitudinally over a varying number of years. See figure A1 for a graphical representation of change in raw scores for each individual/task. The average time span between first and last data points was 6.9 years (std dev. = 3.1 years). The average number of time points was 2.97 (std dev. = 0.77,) with a minimum of 2 and a maximum of 4. The mean slope of verbal fluency was 0.16 std dev. = 1.01) and trail making was −0.03 (std dev. = 0.144). These scores were normalised and combined in order to give a relative composite measure of changes in cognitive performance within the group.

**Figure S1:**
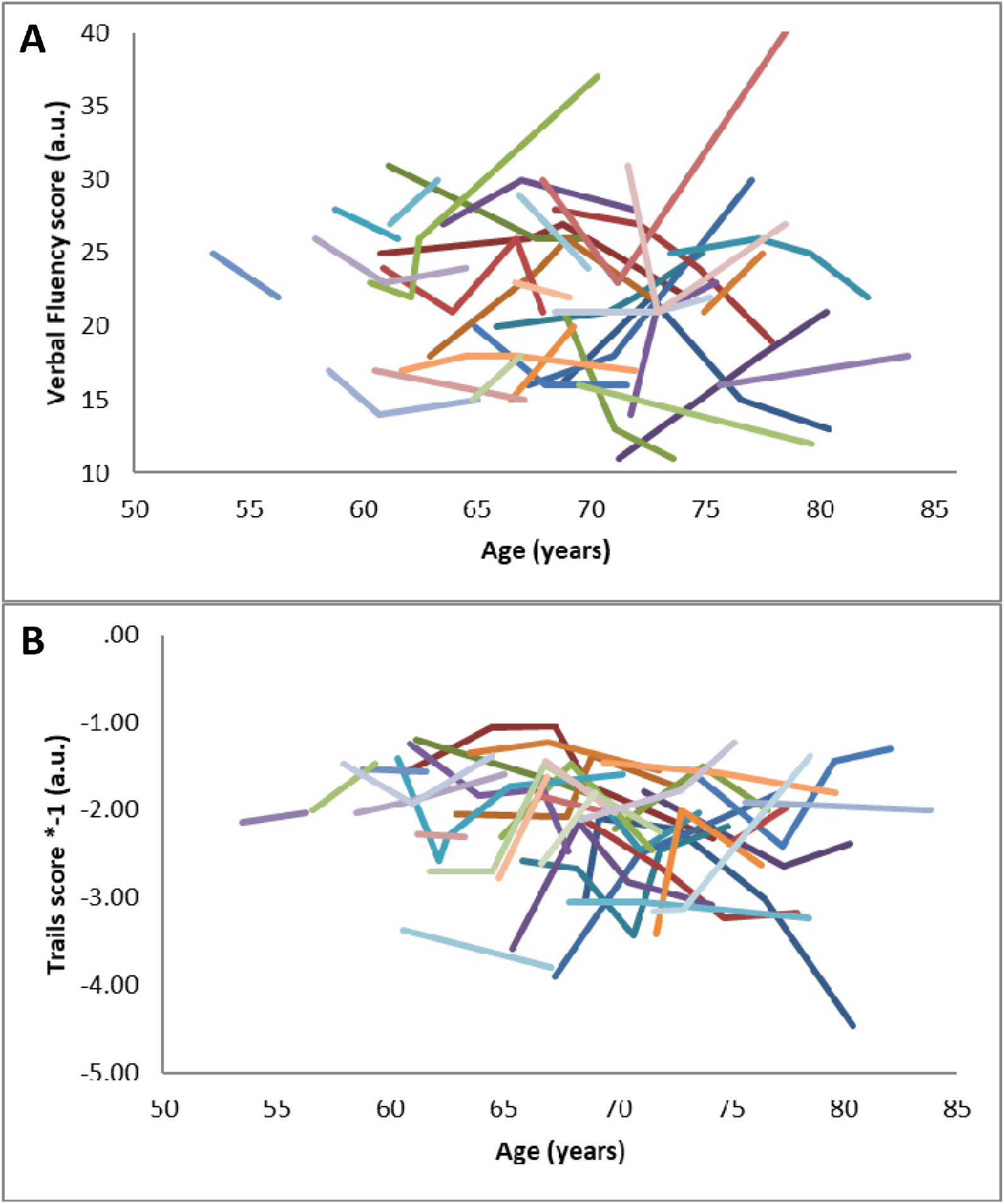
Graphs demonstrating the change in scores for each participant in the verbal fluency category (A) and trails tasks (B).

## Footnotes

1 The full instructions provided during the training with respect to emotion regulation were: “On some trials we will ask you to either INCREASE ("enhance") or DECREASE ("suppress") the intensity of the emotion you are feeling in response to the picture. On other trials we will simply ask you to MAINTAIN ("maintain") the intensity of your emotion. When you hear the instruction "suppress", we would like you to decrease the intensity of the emotion you are feeling when viewing the picture. You may do so by imagining a less negative outcome of the situation in the picture. As you are doing this, please continue to consider the situation to be real. On the other hand, when you hear the instruction "enhance", we would like you to increase the intensity of the emotion you are feeling when you are viewing the picture. You may do this by imagining a worse outcome of the situation in the picture. Finally, when you hear the instruction "maintain", please just maintain the intensity of the emotion you are feeling in response to the picture. You can do this by keeping in mind your initial reaction to the picture. Regardless of the instruction, you should keep focussed on the content of the picture. In other words, please do not think of something unrelated to the picture, and do not try to generate an opposite emotion. For example, we ask you not to think of something comforting, like your home or a good friend, or think to yourself "it’s not real" to counteract any negative emotion you might experience. You may stop following the instruction ("enhance", "suppress", "maintain") as soon as the picture disappears from the screen.”

2 Instead of aggregating scores across the verbal fluency-letter and -category task, we opted to include the category part of the verbal fluency task only, because the letter and category raw scores of the last administration of tests were not correlated (*r* < .1), nor were the longitudinal slopes (*r*(32) = -.21, *p* = .23) in our sample. Dissociations between letter and category fluency with respect to involvement of cortical/frontal areas have also been reported elsewhere (for recent findings with a large N see Vonk et al., 2018).

3 To assess the correlation between age, EF, and rLEF and to ensure that collinearity was not influencing the model, we calculated variance inflation factors (VIFs) for each. All VIFs were sufficiently low (age=1.08, EF=1.32,rLEF=1.29) to allow the assumption that multicollinearity can be safely ignored.

## References

Banks, Sarah J., et al. (2007), Amygdala–frontal connectivity during emotion regulation, Social cognitive and affective neuroscience, 2 (4), 303–12.

Beauregard, M., Levesque, J., and Bourgouin, P. (2001), Neural correlates of conscious self-regulation of emotion, J Neurosci, 21 (18), RC165.

Braunstein, L.M., Gross, J.J., & Ochsner, K.N. (2017). Explicit and implicit emotion regulation: a multi-level framework. Social Cognitive and Affective Neuroscience, 12(10), 1545–1557. https://doi.org/10.1093/scan/nsx096

Buckner, R. L. (2004). Memory and Executive Function in Aging and AD: Multiple Factors that Cause Decline and Reserve Factors that Compensate. Neuron, 44(1), 195–208. https://doi.org/10.1016/j.neuron.2004.09.006

Burgmans, Saartje, et al. (2011), Linking gray matter volume and memory-related activation in the aging brain, Alzheimer’s & Dementia, 7 (4, Supplement), S419.

Calhoun, V. (2018), Data-driven approaches for identifying links between brain structure and function in health and disease, Dialogues Clin Neurosci, 20(2), 87–99.

Carmichael, S. T. and Price, J. L. (1995), Limbic connections of the orbital and medial prefrontal cortex in macaque monkeys, J Comp Neurol, 363 (4), 615–41.

Desikan, R. S., Ségonne, F., Fischl, B., Quinn, B. T., Dickerson, B. C., Blacker, D., … Killiany, R. J. (2006). An automated labeling system for subdividing the human cerebral cortex on MRI scans into gyral based regions of interest. NeuroImage, 31(3), 968–980. https://doi.org/10.1016/j.neuroimage.2006.01.021

Eippert, F., et al. (2007), Regulation of emotional responses elicited by threat-related stimuli, Hum Brain Mapp, 28 (5), 409–23.

Fleischman, D. A., et al. (2014), Gray-matter macrostructure in cognitively healthy older persons: associations with age and cognition, Brain Structure and Function, 219 (6), 2029–49.

Giorgio, Antonio, et al. (2010), Age-related changes in grey and white matter structure throughout adulthood, NeuroImage, 51 (3), 943–51.

Goldin, P. R., et al. (2008), The neural bases of emotion regulation: reappraisal and suppression of negative emotion, Biol Psychiatry, 63 (6), 577–86.

Gunning-Dixon, F. M., et al. (2009), Aging of cerebral white matter: a review of MRI findings, Int J Geriatr Psychiatry, 24 (2), 109–17.

Gyurak, A., Goodkind, M., Kramer, J., Miller, B., & Levenson, R. (2011). Executive functions and the down-regulation and up-regulation of emotion. Cognition & Emotion, 1–16. https://doi.org/10.1080/02699931.2011.557291

Gyurak, A., Goodkind, M. S., Madan, A., Kramer, J. H., Miller, B. L., & Levenson, R. W. (2009). Do tests of executive functioning predict ability to downregulate emotions spontaneously and when instructed to suppress? Cognitive, Affective, & Behavioral Neuroscience, 9(2), 144–152. https://doi.org/10.3758/CABN.9.2.144

Hakun, J. G., Zhu, Z., Brown, C. A., Johnson, N. F., & Gold, B. T. (2015). Longitudinal alterations to brain function, structure, and cognitive performance in healthy older adults: A fMRI-DTI study. Neuropsychologia, 71, 225–235. https://doi.org/10.1016/j.neuropsychologia.2015.04.008

Jenkinson, M., Beckmann, C. F., Behrens, T. E. J., Woolrich, M. W., & Smith, S. M. (2012). FSL. NeuroImage, 62(2), 782–790. https://doi.org/10.1016/j.neuroimage.2011.09.015

Johnstone, T., et al. (2007), Failure to regulate: counterproductive recruitment of top-down prefrontal-subcortical circuitry in major depression, J Neurosci, 27 (33), 8877–84.

Kalisch, R., et al. (2006), Neural correlates of self-distraction from anxiety and a process model of cognitive emotion regulation, J Cogn Neurosci, 18 (8), 1266–76.

Kalpouzos, G., Persson, J., & Nyberg, L. (2012). Local brain atrophy accounts for functional activity differences in normal aging. Neurobiology of Aging, 33(3), 623.e1–623.e13. https://doi.org/10.1016/j.neurobiolaging.2011.02.021

Kim, S. H. and Hamann, S. (2007), Neural correlates of positive and negative emotion regulation, J Cogn Neurosci, 19 (5), 776–98.

Lang, Peter, Bradley, Margaret, and Cuthbert, B. N. (2008), International affective picture system (IAPS): Affective ratings of pictures and instruction manual.

Manard, M., et al. (2016), Relationship between grey matter integrity and executive abilities in aging, Brain Research, 1642, 562–80.

Mather, M. (2012). The emotion paradox in the aging brain. Annals of the New York Academy of Sciences, 1251(1), 33–49. https://doi.org/10.1111/j.17496632.2012.06471.x

Mazziotta, J. C., Toga, A. W., Evans, A., Fox, P., & Lancaster, J. (1995). A Probabilistic Atlas of the Human Brain: Theory and Rationale for Its Development: The International Consortium for Brain Mapping (ICBM). NeuroImage, 2(2, Part A), 89–101. https://doi.org/10.1006/nimg.1995.1012

Miller, E. (1984). Verbal fluency as a function of a measure of verbal intelligence and in relation to different types of cerebral pathology. British Journal of Clinical Psychology, 23(1), 53–57. https://doi.org/10.1111/j.2044-8260.1984.tb00626.x

Morawetz, C., Bode, S., Derntl, B., & Heekeren, H. R. (2017). The effect of strategies, goals and stimulus material on the neural mechanisms of emotion regulation: A meta-analysis of fMRI studies. Neuroscience & Biobehavioral Reviews, 72, 111–128. https://doi.org/10.1016/j.neubiorev.2016.11.014

Morcom, A. M. and Johnson, W. (2015), Neural reorganization and compensation in aging, J Cogn Neurosci, 27 (7), 1275–85.

Mori, S., Wakana, S., Zijl, P. C. M. van, & Nagae-Poetscher, L. M. (2005). MRI Atlas of Human White Matter. Elsevier.

Opitz, P. C., Lee, I. A., Gross, J. J., & Urry, H. L. (2014). Fluid cognitive ability is a resource for successful emotion regulation in older and younger adults. Frontiers in Psychology, 5. https://doi.org/10.3389/fpsyg.2014.00609

Ochsner, Kevin N., et al. (2002), Rethinking Feelings: An fMRI Study of the Cognitive Regulation of Emotion, Journal of Cognitive Neuroscience, 14 (8), 1215–29.

Ochsner, K.N., et al. (2004), For better or for worse: neural systems supporting the cognitive down- and up-regulation of negative emotion, Neuroimage, 23 (2), 483–99.

Ochsner K.N., Gross J.J. (2005). The cognitive control of emotion. Trends in Cognitive Science, 9, 242–9.

Ochsner K.N., Silvers J., Buhle J.T. (2012). Functional imaging studies of emotion regulation: a synthetic review and evolving model of the cognitive control of emotion. Annals of the New York Academy of Sciences, 1251, E1–24. doi:10.1111/j.1749-6632.2012.06751.x.

Ongur, D. and Price, J. L. (2000), The organization of networks within the orbital and medial prefrontal cortex of rats, monkeys and humans, Cereb Cortex, 10 (3), 206–19.

Opitz, PC, et al. (2012), Prefrontal mediation of age differences in cognitive reappraisal, Neurobiology of Aging, 33 (4), 645–55.

Park, D. C. and Reuter-Lorenz, P. (2009), The adaptive brain: aging and neurocognitive scaffolding, Annu Rev Psychol, 60, 173–96.

Phan, K. L., et al. (2005), Neural substrates for voluntary suppression of negative affect: a functional magnetic resonance imaging study, Biol Psychiatry, 57 (3), 210–9.

Phillips M.L., Ladouceur C.D., Drevets W.C. (2008). A neural model of voluntary and automatic emotion regulation: implications for understanding the pathophysiology and neurodevelopment of bipolar disorder. Molecular Psychiatry, 13, 829–57.

Raz, N., & Rodrigue, K. M. (2006). Differential aging of the brain: patterns, cognitive correlates and modifiers. Neuroscience & Biobehavioral Reviews, 30(6), 730–748.

Reitan, R. M., & Wolfson, D. (1985). The Halstead-Reitan neuropsychological test battery: Theory and clinical interpretation. (Vol. 4). Tuscon, AZ:Neurospychological Press.

Sala-Llonch, R., Bartrés-Faz, D., Junqué, C. (201). Reorganization of brain networks in aging: a review of functional connectivity studies. Front Psychol. 21(6)663. doi:10.3389/fpsyg.2015.00663

Salthouse, T. A. (2011), Neuroanatomical substrates of age-related cognitive decline, Psychological Bulletin, 137 (5), 753–84.

Scheibe, S. and Carstensen, L. L. (2010), Emotional aging: recent findings and future trends, J Gerontol B Psychol Sci Soc Sci, 65B (2), 135–44.

Smith, S. M., Jenkinson, M., Woolrich, M. W., Beckmann, C. F., Behrens, T. E. J., Johansen-Berg, H., … Matthews, P. M. (2004). Advances in functional and structural MR image analysis and implementation as FSL. NeuroImage, 23, Supplement 1, S208–S219. https://doi.org/10.1016/j.neuroimage.2004.07.051

Swartz, J. R., Carrasco, M., Wiggins, J. L., Thomason, M. E., & Monk, C. S. (2014). Age-related changes in the structure and function of prefrontal cortex– amygdala circuitry in children and adolescents: A multi-modal imaging approach. NeuroImage, 86, 212–220. https://doi.org/10.1016/j.neuroimage.2013.08.018

Teipel, S. J., Bokde, A. L.., Born, C., Meindl, T., Reiser, M., Möller, H. J., & Hampel, H. (2007). Morphological substrate of face matching in healthy ageing and mild cognitive impairment: a combined MRI-fMRI study. Brain, 130(7), 1745.

Urry, Heather L, et al. (2006), Amygdala and ventromedial prefrontal cortex are inversely coupled during regulation of negative affect and predict the diurnal pattern of cortisol secretion among older adults, The Journal of Neuroscience, 26 (16), 4415–25.

Urry, Heather L., et al. (2009), Individual differences in some (but not all) medial prefrontal regions reflect cognitive demand while regulating unpleasant emotion, NeuroImage, 47 (3), 852–63.

Urry, H. L., & Gross, J. J. (2010). Emotion Regulation in Older Age. Current Directions in Psychological Science, 19(6), 352–357. https://doi.org/10.1177/0963721410388395

Van Petten, C., Plante, E., Davidson, P. S. R., Kuo, T. Y., Bajuscak, L., & Glisky, E. L. (2004). Memory and executive function in older adults: relationships with temporal and prefrontal gray matter volumes and white matter hyperintensities. Neuropsychologia, 42(10), 1313–1335. https://doi.org/10.1016/j.neuropsychologia.2004.02.009

van Reekum, C. M, Johnstone, T., Urry, H. L., Thurow, M. E., Schaefer, H. S., Alexander, A. L., & Davidson, R. J. (2007). Gaze fixations predict brain activation during the voluntary regulation of picture-induced negative affect. Neuroimage, 36(3), 1041–1055.

van Reekum, Carien M., Schaefer, S. M., Lapate, R. C., Norris, C. J., Tun, P. A., Lachman, M. E., … Davidson, R. J. (2018). Aging is associated with a prefrontal lateral-medial shift during picture-induced negative affect. Social Cognitive and Affective Neuroscience, 13(2), 156–163. https://doi.org/10.1093/scan/nsx144

Vonk, J. M. J., Rizvi, B., Lao, P. J., Budge, M., Manly, J. J., Mayeux, R., & Brickman, A. M. (n.d.). Letter and Category Fluency Performance Correlates with Distinct Patterns of Cortical Thickness in Older Adults. Cerebral Cortex. https://doi.org/10.1093/cercor/bhy138

Wager, T. D., Davidson, M. L., Hughes, B. L., Lindquist, M. A., & Ochsner, K. N. (2008). Prefrontal-Subcortical Pathways Mediating Successful Emotion Regulation. Neuron, 59(6), 1037–1050. https://doi.org/10.1016/j.neuron.2008.09.006

Winecoff, A, et al. (2011), Cognitive and neural contributors to emotion regulation in aging, Social Cognitive and Affective Neuroscience, 6 (2), 165–76.

Ziegler, D. A., Piguet, O., Salat, D. H., Prince, K., Connally, E., & Corkin, S. (2010). Cognition in healthy aging is related to regional white matter integrity, but not cortical thickness. Neurobiology of Aging, 31(11), 1912–1926. https://doi.org/10.1016/j.neurobiolaging.2008.10.015

